# Spatially Resolved Single-cell Translatomics at Molecular Resolution

**DOI:** 10.1101/2022.09.27.509605

**Authors:** Hu Zeng, Jiahao Huang, Jingyi Ren, Connie Kangni Wang, Zefang Tang, Yiming Zhou, Abhishek Aditham, Hailing Shi, Xin Sui, Xiao Wang

**Affiliations:** Department of Chemistry, Massachusetts Institute of Technology, Cambridge, MA 02139, USA; Broad Institute of MIT and Harvard, Cambridge, MA 02142, USA; Center for Psychiatric Research, Broad Institute of MIT and Harvard, Cambridge, MA 02142, USA; Department of Biological Engineering, Massachusetts Institute of Technology, Cambridge, MA 02139, USA

## Abstract

The precise control of mRNA translation is a crucial step in post-transcriptional gene regulation of cellular physiology. However, it remains a major challenge to systematically study mRNA translation at the transcriptomic scale with spatial and single-cell resolutions. Here, we report the development of RIBOmap, a three-dimensional (3D) *in situ* profiling method to detect mRNA translation for thousands of genes simultaneously in intact cells and tissues. By applying RIBOmap to 981 genes in HeLa cells, we revealed remarkable dependency of translation on cell-cycle stages and subcellular localization. Furthermore, we profiled single-cell translatomes of 5,413 genes in the adult mouse brain tissue with a spatial cell atlas of 62,753 cells. This spatial translatome profiling detected widespread patterns of localized translation in neuronal and glial cells in intact brain tissue networks. Together, RIBOmap presents the first spatially resolved single-cell translatomics technology, accelerating our understanding of protein synthesis in the context of subcellular architecture, cell types, and tissue anatomy.

## Main Text

Cellular proteins are final products of gene expression that execute cellular functions. Measuring the genome-wide protein synthesis patterns with single-cell and spatial resolution is a paramount goal that can transform our understanding of translational regulation in heterogeneous cell types and states. While large-scale single-cell and spatially resolved proteome profiling remains challenging (*1*), past research has been focused on mapping mRNA levels to infer the corresponding protein abundances in single cells. However, numerous studies have concluded poor correlations between mRNA and protein levels (*2*–*5*) as gene regulation is achieved at both transcription and translation levels. Moreover, many genes undergo signal-dependent and subcellular-localized translation (*6*), adding another layer of complexity in regulating when and where different proteins are produced. Therefore, we need scalable single-cell and spatially resolved profiling of protein synthesis for a comprehensive understanding of gene expression and translational regulation.

Existing bulk and single-cell ribosome profiling methods have enabled us to analyze protein translation at the transcriptome scale (*7*–*12*). However, they cannot preserve the spatial information of subcellular structure, cell morphology, or tissue organization, limiting our ability to decipher the regulation of protein synthesis in its biological context. In contrast, imaging-based methods that can trace mRNA translation with their physical coordinates (*13*–*19*) are limited to one gene at a time. Therefore, it remains challenging to achieve highly multiplexed spatial ribosome profiling in single cells with subcellular resolution. To fill this gap, here we develop a new 3D *in situ* ribosome-bound mRNA mapping method (RIBOmap) for highly multiplexed charting of protein synthesis with single-cell and subcellular resolutions. We profiled the translation of 981 genes in cultured cells with RIBOmap and developed computational pipelines to analyze different subcellular patterning of localized translation. We further applied RIBOmap to intact mouse brain tissues, profiling 5,413 genes at subcellular resolution, which revealed spatial patterns of protein synthesis across various cell types and tissue regions.

RIBOmap is built upon a targeted-sequencing strategy where a unique design of tri-probes selectively detects and amplifies ribosome-bound mRNAs, followed by hydrogel-tissue embedding and highly multiplexed *in situ* sequencing readout (*20*) (Fig. 1A). The RIBOmap tri-probe set includes: (1) a splint DNA probe that hybridizes to ribosomal RNAs (rRNAs) and serves as the splint to circularize nearby padlock probes; (2) a padlock probe that targets specific mRNA species of interest and encodes a gene-unique identifier; (3) a primer that targets the adjacent site of the padlock probe on the same mRNA and serves as the primer to enable amplification of the circularized padlock probes through rolling circle amplification (RCA), resulting in a DNA nanoball (amplicon). Importantly, the gene-specific pair of primer-padlock probes excludes false-positive signals arising from the nonspecific hybridization of a single probe (*20*). Together, only when all three probes are present in proximity, the signal of RNAs can be amplified to DNA amplicons (Fig. 1, B and C). The DNA amplicons are subsequently embedded *in situ* in a polyacrylamide hydrogel matrix using tissue-hydrogel conjugation chemistry (*20*). The gene-unique barcodes in the DNA amplicons are then decoded through *in situ* sequencing with error-reduction by dynamic annealing and ligation (SEDAL, Fig. 1A) (*20*).

**Fig. 1.**
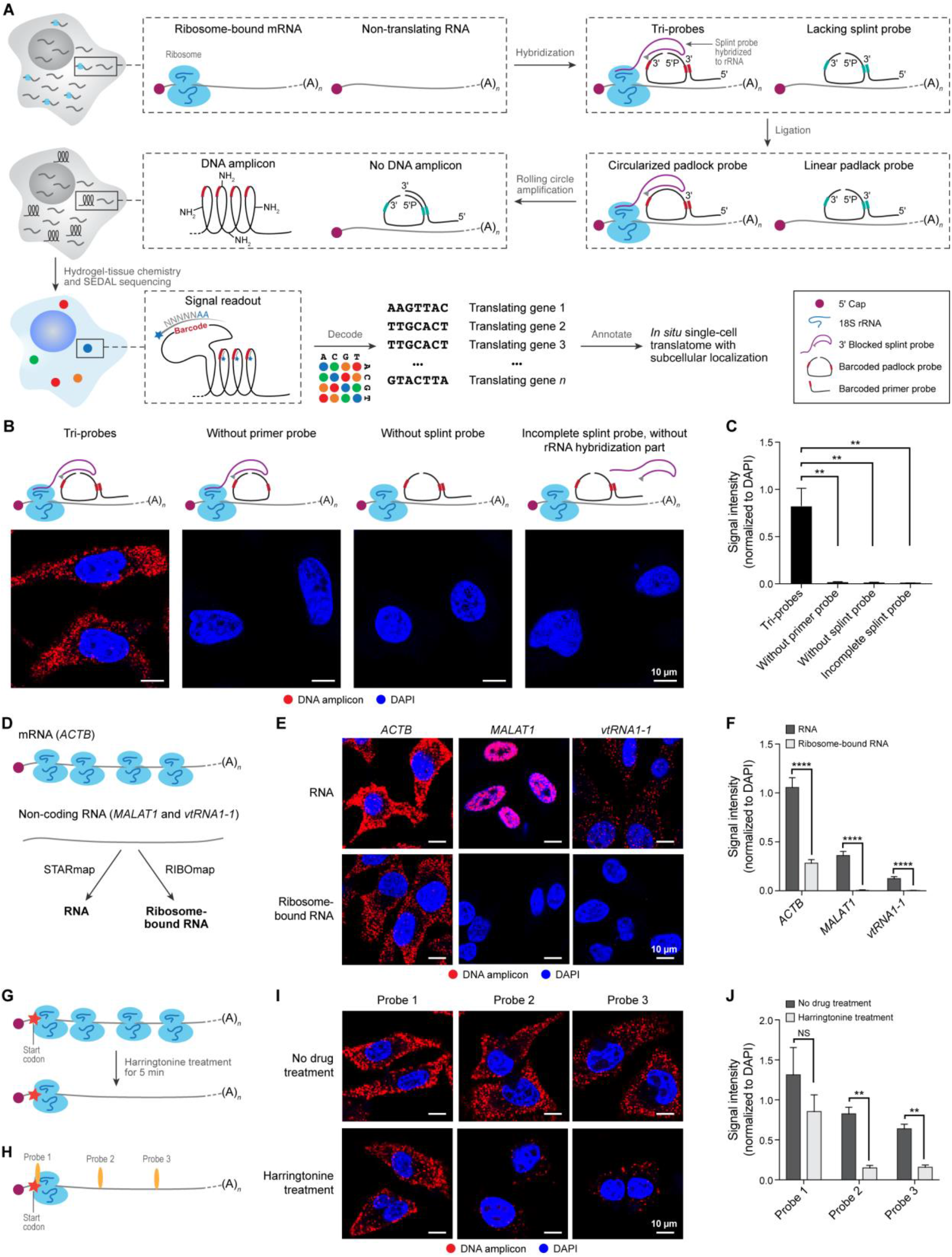
RIBOmap for *in situ* profiling of mRNA translation at subcellular resolution. (**A**) Schematic of RIBOmap. After the sample (cell line or tissue sample) is prepared, a pair of barcoded padlock probes and primers are hybridized to a targeted intracellular RNA, and the splint probe is hybridized to 18S rRNA of ribosomes. The splint probe is used as a template for the proximity ligation to circularize the padlock probe. The splint probe is designed with 3’ Inverted dT modifications to block its extension during RCA. The intact padlock probe can then be amplified to generate amine-modified DNA amplicons *in situ*. Next, these DNA amplicons are copolymerized into hydrogel via tissue-hydrogel chemistry for *in situ* sequencing. The gene-specific identifier sequence (red) in the cDNA amplicons can then be read out through cyclic *in situ* sequencing with error reduction by dynamic annealing and ligation (SEDAL). (**B)** Tri-probe strategy. (Left) Fluorescent images of tri-probe condition show the RIBOmap signal of *ACTB* mRNA in HeLa cells. (Middle) Fluorescent images of negative control samples without the primer or splint probe show minimum DNA amplicon signal. (Right) Fluorescent images of the control sample using splint probes without the rRNA hybridization sequence show minimal DNA amplicon signal. (**C**) Quantification of the DNA amplicon signal intensity normalized to the DAPI signal in panel B. Error bars, standard deviation. *n* = 3 images per condition. Student’s t-test, \*\**P* < 0.01. (**D**) Schematic representation of RIBOmap signal verification by targeting mRNA (*ACTB*) and non-coding RNA (*MALAT 1* and *vtRNA1-1*). (**E**) Fluorescent images show RNA detection results by STARmap and translating RNA detection results by RIBOmap. (**F**) Quantification of the DNA amplicon signal intensity that was normalized to the DAPI signal is shown in panel E. Error bars, standard deviation. *n* = 5 images per condition. Student’s t-test, \*\*\*\**P* < 0.0001. (**G**) Schematic of translational regulation by Harringtonine. (**H**) Three pairs of SNAIL probes targeting different sites of *ACTB* mRNA. Probe pair 1 is 16 nt from the start codon, while Probe pair 2 and Probe pair 3 are 115 nt and 405 nt downstream of the start codon in the CDS region, respectively. Fluorescent images show the RIBOmap signal of 3 sets of probes targeted to different regions of *ACTB* mRNA in HeLa cells before and after Harringtonine treatment. (**J**) Quantification of the RIBOmap signal intensity that was normalized to the DAPI signal is shown in panel I. Error bars, standard deviation. *n* = 3 images per condition. Student’s t-test, \*\**P* < 0.01.

To further validate the necessity of targeting the rRNA to detect ribosome-bound mRNAs, we also tested the alternative strategy for RIBOmap workflow that utilizes primary antibodies to target ribosomal proteins, which can be subsequently detected by splint-conjugated Protein A/G for tri-probe amplification (fig. S1A). We evaluated this strategy using anti-RPS3 (small 40S subunit ribosomal protein 3) antibody and anti-RPL4 (large 60S subunit ribosome protein 4) antibody (fig. S1, B to E). However, the signal-to-noise ratio (SNR) is in the range of 14-17, much lower than that of the rRNA-targeting strategy (SNR = 93±17, fig. S1F). Thus, we finalized with the rRNA-targeting strategy in the following studies.

A key factor for the success of RIBOmap is whether the RIBOmap tri-probe amplification is specific for ribosome-bound mRNAs. To test this, we first used non-translating non-coding RNAs (nuclear-localized *MALAT1* RNA and cytoplasmic localized *vtRNA1-1* RNA) as negative controls. We performed RIBOmap to detect ribosome-bound RNAs and previously reported STARmap RNA imaging (*20*) for *ACTB* mRNA and negative control (non-coding) RNAs in HeLa cells (Fig. 1D). As expected, we found that RIBOmap only detected *ACTB* mRNAs while STARmap detected both *ACTB* mRNA and non-coding RNAs (Fig. 1, E and F). Then, we validated the sub-transcript specificity of RIBOmap in detecting ribosome-loading sites, by using Harringtonine, a translation inhibitor that traps the translation-initiating ribosomes at the start codon and allows the elongating ribosomes to run off the transcript (Fig. 1G). Three tri-probe sets were designed to target sub-transcript regions at different distances from the *ACTB* start codon (−16, 115, 405 nt) (Fig. 1H). After 5 min of Harringtonine treatment, the *ACTB* mRNA showed a significant decrease in RIBOmap signal intensity when detected by the probes targeting 115 or 405 nt downstream of the start codon (*p* = 7.5 × 10^−5^ and 4.7 × 10^−5^ respectively, student’s *t*-test), while signals generated by the probes targeting the -16 nt from the start codon showed little decrease (Fig. 1, I and J). Altogether, the results demonstrated the specificity of RIBOmap in detecting ribosome-bound mRNAs.

After benchmarking the SNR and specificity of RIBOmap, we set out to uncover cell-cycle dependent and localized mRNA translation at subcellular resolution in single cells by performing a highly multiplexed 981-gene RIBOmap experiment in HeLa cells (Fig. 2A). The 981 genes represent a curated list composed of cell-cycle gene markers, genes with diverse subcellular RNA patterns, and genes of varying RNA stabilities (*21*–*24*). We designed a multi-modal RIBOmap imaging experiment that further incorporated the information on cell-cycle stages and subcellular organelles: (1) the cell-cycle phases of each cell were captured by the fluorescent ubiquitination-based cell cycle indicators (FUCCI) (*25, 26*); (2) ribosome-bound mRNAs (981 genes) were *in situ* sequenced after FUCCI imaging (*20*); (3) finally, nuclei, endoplasmic reticulum, and cell shapes were stained and imaged (Fig. 2B). To compare the single-cell spatial translatome versus the spatial transcriptome, we also conducted a paired total RNA *in situ* sequencing of the same 981 genes using STARmap (with identical RNA hybridization sequences) for the same batch of HeLa FUCCI cells (Fig. 2A). In total, we sequenced 1,813 cells by RIBOmap and 1,757 cells by STARmap (fig. S2A).

**Fig. 2.**
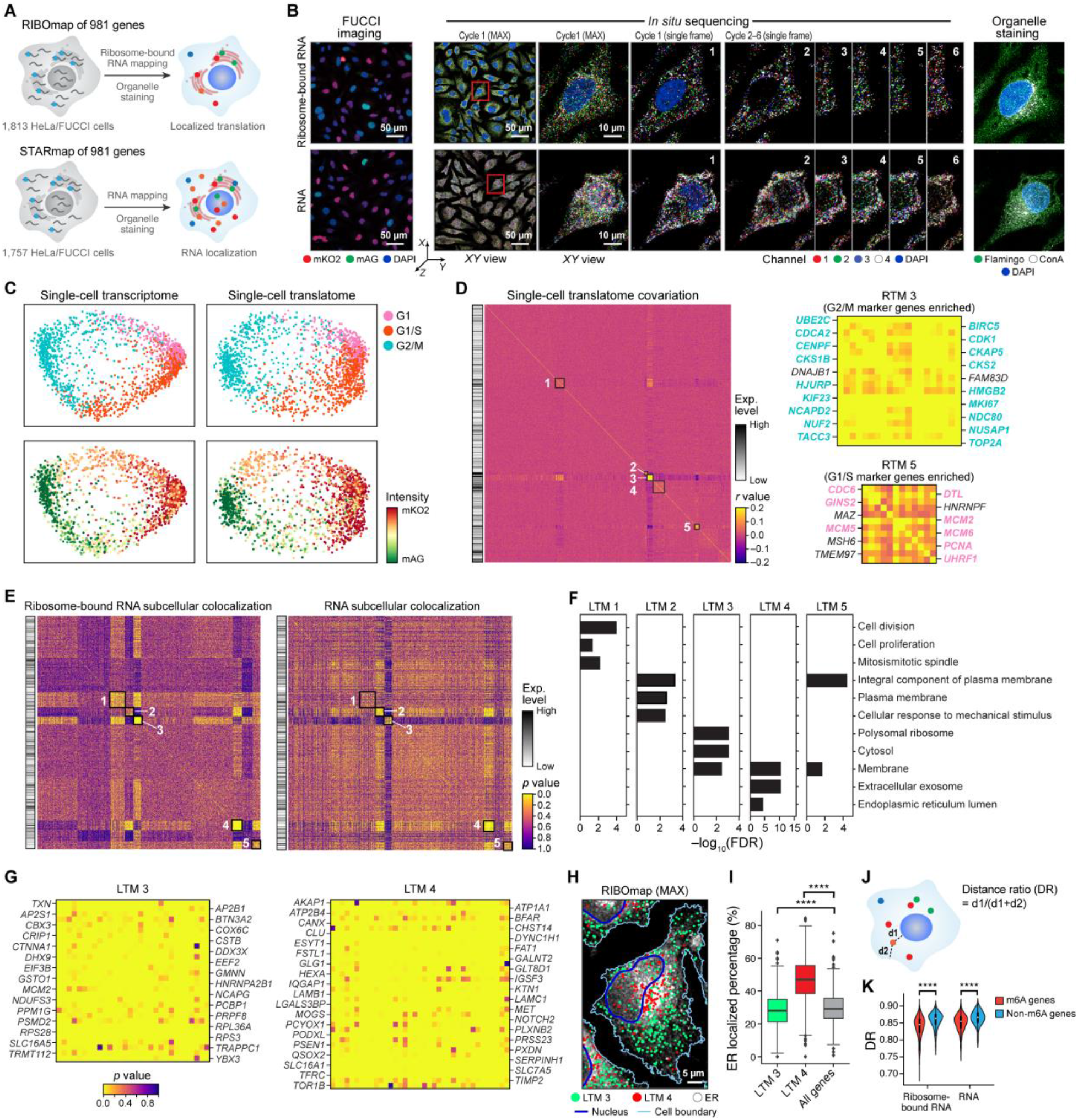
RIBOmap simultaneously measures the subcellular translation of 981 genes in human HeLa cells. (**A**) Schematic of RIBOmap detection in HeLa FUCCI cells to measure localized mRNA translation. RIBOmap reads out ribosome-bound mRNA signals and STARmap detects all the targeted RNA. Samples were also subjected to organelle staining to visualize nuclei, ER and cell morphology. (**B**) Representative images showing the sequential mapping of FUCCI fluorescence signal, cDNA amplicons, and organelle staining in the same HeLa cell sample. Left, the FUCCI fluorescence imaging results of the cells for RIBOmap (upper) and STARmap (bottom). The fluorescent signals indicate the cell cycle stage of HeLa cells. Middle, representative images showing the maximum-intensity projections (MAX) of the first sequencing cycle for 981 genes with zoom-in views of a representative cell and single-frame views of the representative cell across 6 sequencing cycles. Right, magnified view of organelle staining of a representative cell. White, Alexa Fluor 594 Conjugate of Concanavalin A (ConA) staining of ER; green, Flamingo fluorescent staining of cell morphology; blue: DAPI staining of nucleus. (**C**) Diffusion map embeddings of cell-cycle stage clusters determined using the single-cell expression profile of 41 cell-cycle marker genes for STARmap (upper left) and RIBOmap (upper right) measurements along with corresponding protein fluorescence profiles of the monomeric Kusabira-Orange 2 (mKO2)–hCDT1 and monomeric Azami-Green (mAG)–hGEN FUCCI cell-cycle markers (STARmap, bottom left; RIBOmap, bottom right). (**D**) Single-cell translatome covariation matrix showing the pairwise Pearson’s correlation coefficients of the cell-to-cell variation, shown together with their averaged expression levels. Five strongly correlating blocks of co-regulated translation modules (RTMs) are indicated by the gray boxes in the matrix, with RTMs 3 and 5 enlarged on the right. Cell cycle markers are highlighted in the enlarged matrix. (**E**) Matrix of the pairwise colocalization p-values describing the degree to which the reads of two genes tend to co-exist in a sphere of 3-µm radius in the same cell in RIBOmap results (left) and STARmap results (right), shown together with the expression levels of these genes. The STARmap matrix uses the same order of genes as the RIBOmap matrix. Five strongly correlating blocks of co-localized translation modules (LTMs) are indicated by the gray boxes in the matrix. (**F**) Bar plots visualizing the most significantly enriched Gene Ontology (GO) terms (maximum 3) in each of the five gene modules. (**G**) Enlarged gene matrix of LTMs 3 and 4. (**H**) A representative cell image showing the spatial distribution of RIBOmap signals of LTMs 3 and 4 genes, overlaid on the ER and cell boundary. The images are MAXs of the gene dots and ER signals. (**I**) Quantification of the ER-localized percentage of genes of LTMs 3 and 4 versus all the detected genes. Wilcoxon signed-rank test, \*\*\*\**P* < 0.0001. (**J**) Diagram showing the calculation of spatial parameter Distance Ratio (DR). (**K**) Violin plots for the ribosome-bound RNA DR and RNA DR values for m^6^A genes and non-m^6^A genes. Wilcoxon signed-rank test, \*\*\*\**P* < 0.0001.

Next, we evaluated whether RIBOmap can decipher cell cycle-dependent mRNA translation. To this end, we treated the single-cell protein fluorescence profiles of the FUCCI as the ground truth of cell-cycle phases and compared them with the single-cell profiles of RIBOmap and STARmap measurements. Specifically, known cell-cycle gene markers (*21*) from the single-cell translatomic profiles of RIBOmap and the single-cell transcriptional profiles of STARmap were used to embed single cells on the diffusion maps to delineate the trajectory of cell-cycle progression (Fig. 2C and fig. S2, B and C). The results showed that the G1, G1/S, and G2/M cell-cycle phases identified from both RIBOmap and STARmap datasets agreed with the expected cell-cycle-dependent patterns of FUCCI protein fluorescence (Fig. 2C), demonstrating the accuracy of RIBOmap in delineating cell states.

We then aimed to identify potential translationally co-regulated gene modules among single cells by covariation analysis. We calculated the pairwise correlations for all gene pairs based on single-cell translatomic expression and used hierarchical clustering to identify genes with high correlation coefficients across single cells. We identified five co-regulated translation modules (RTMs) with substantial intra-module correlation, each with enrichment of distinct functional pathways (RTMs 1-5, Fig. 2D and fig. S3, A to D). RTM 3 and RTM 5 are enriched for G2/M and G1/S cell-cycle marker genes, respectively (Fig. 2E), and these two gene modules are negatively correlated (fig. S3E). This result provided the single-cell level evidence corroborating the previous finding that these cell-cycle marker genes are co-regulated during cell-cycle progression for synchronized execution of their gene functions (*27*). Moreover, RTM 2 contains genes encoding protein translation machineries such as ribosomal proteins (*RPS3, RPS28*, and *RPL37*) and translational factors (*EEF2, EIF3B*, and *ETF1*) (fig. S3C); these genes show a positive correlation with RTM 5 (G1/S marker genes) and a negative correlation with RTM 3 (G2/M marker genes) (fig. S3F). This discovery implies that the protein translation machinery is upregulated in the G1/S phase to support the demand for protein products during the physical expansion of cells and the replication of subcellular organelles.

Subcellular localized translation has great implications for signal transduction and proteome organization (*6*). Leveraging the high spatial resolution of the dataset, we explored whether RIBOmap could detect subcellular patterns of translation. To identify translating genes that show a tendency to co-localize in the subcellular space, we generated the nearest neighbor profile for each RNA in RIBOmap and evaluated the significance of gene co-localization by comparing against a randomized control. With subsequent hierarchical clustering of the gene co-localization matrix, we identified five co-localized translation modules (LTMs) with highly correlated subcellular spatial organizations and distinct functional enrichment (LTMs 1-5, Fig. 2, E to G and fig. S4, A to C). Among them, LTMs 2, 4, and 5 are functionally enriched with membrane protein and secretion pathway (Fig. 2, F and G and fig. S4 A and B), and physically co-localize with ER staining (Fig. 2, H and I and fig. S3, D and E), suggesting they are ER-translated genes. In contrast, LTMs 1 and 3 are enriched for genes encoding large protein complexes of mitotic spindle (*e*.*g*., *CDK1, NUSAP1, KIF23, SPAG5, TACC3, FAM83D*) and translation machinery (*e*.*g*., *RPS3, RPS28, RPL36A, EEF2, DHX9*), respectively (Fig. 2, F and G and fig. S4, A and C). Such observation indicates the subunits of large protein complexes may be synthesized in spatial proximity for efficient assembly. Finally, we explored if the co-regulated genes tended to be co-localized for translation. To this end, we plotted the co-localization *p*-values from the subcellular colocalization analysis into the single-cell covariation gene matrix and found that the five RTMs also show strong subcellular colocalization (fig. S4F). These results may imply that functionally related gene groups can be co-regulated through subcellular co-localized translation, possibly through shared regulatory RNA elements or protein-protein interactions of the nascent proteins.

One mechanism to coordinate the translation of mRNA is post-transcriptional RNA modifications (*28*). For example, *N*^6^-methyladenosine modification (m^6^A) is a critical post-transcriptional RNA modification that marks short-lived RNAs and regulates RNA translation (*23, 28*). We hypothesized that the shorter lifetime of m^6^A-modified RNA may lead to a shorter trafficking time of mRNA in the cytosol, and consequently influence the loci of mRNA translation. To test this, we quantified the translation distribution of each gene with a distance-ratio (DR) metric, which estimated the relative position of each amplicon between nuclear and cytoplasmic membranes (Fig. 2J). Notably, RIBOmap illustrated that the DR values of m^6^A genes were significantly lower than those of non-m^6^A genes (Fig. 2K), indicating that m^6^A-modified RNAs are indeed physically translated closer to the nuclear membrane. Together, RIBOmap enables integrative analysis of spatial translatomics at subcellular levels, offering an unprecedented approach for future studies on RNA processing and translation.

With the success of applying RIBOmap in cultured cells, we took the next step to apply RIBOmap to mouse brain tissue slices for revealing single-cell translatomic profiles in intact tissues. Here, we mapped a targeted gene list of 5,413 genes curated from previously published single-cell sequencing studies of mouse cell atlas (*29*–*35*). Through nine rounds of *in situ* sequencing, we imaged a 10 µm-thick coronal section of the mouse left hemisphere (62,753 cells) containing multiple brain regions at 90 nm × 90 nm × 300 nm voxel size (Fig. 3, A and B). We benchmarked tissue RIBOmap data by comparing the spatial translation pattern of well-known cell-type marker genes (e.g., *Slc17a7, Gad2*, and *Pcp4*) with *in situ* hybridization (ISH) images of the corresponding genes from the Allen Brain database (*36*) (Fig. 3C and fig. S5A), where the comparison showed consistent spatial patterns. Notably, we observed that RIBOmap reads of specific genes correlated better with the protein signals in spatial patterns than the RNA signals (ISH) in specific brain regions, such as *Sst* in CA3 and *Nefl* in CA1 of the hippocampus (fig. S5, B and C). This result suggests that RIBOmap has better SNR than ISH or the translatome has better correlation with the proteome than the transcriptome (*7, 11, 12*).

**Fig. 3.**
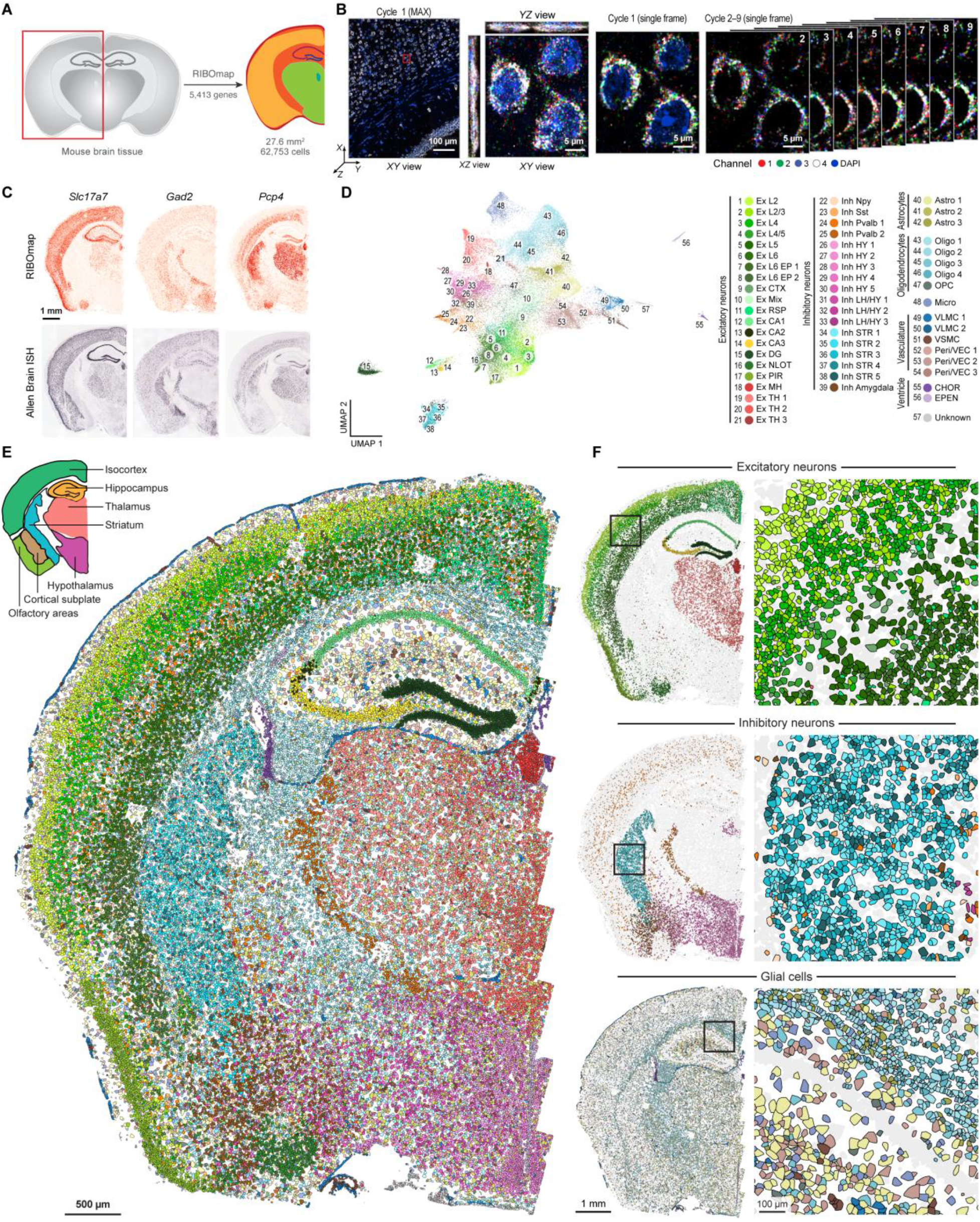
Spatially single-cell translatomic profiling of 5,413 genes in the mouse brain. (**A**) Diagram of the imaged mouse coronal hemibrain region (red box) for RIBOmap. (**B**) Representative images showing the measurements of localized translation of 5,413 genes by RIBOmap in a mouse coronal hemibrain slice. (Left) Maximum-intensity projection of the first sequencing round of 5,413 genes, showing all five channels simultaneously. Red square, zoom region. (Middle) Magnified view of three cells showing the maximum-intensity projection (MAX) view of the first sequencing round. (Right) Magnified view of three cells showing the spatial arrangement of amplicons in a single z-frame across nine sequencing rounds. (**C**) RIBOmap images of three cell-type marker genes in comparison with the Allen Brain ISH images (*36*) showing the expression patterns of the corresponding genes. (**D**) Uniform Manifold Approximation and Projection (UMAP) plot visualization of translational profiles of 62,753 cells collected from mouse coronal hemibrain. 57 cell types were identified using Leiden clustering. (**E**) Representative spatial cell-type atlas in the imaged coronal hemibrain region using the same color code as in (D). (**F**) The spatial cell map of excitatory neuron subtypes (upper), inhibitory neuron subtypes (middle), and glial cell subtypes (bottom) in the imaged coronal hemibrain region with magnified views of spatial cell-type maps.

Encouraged by the benchmark results, we then subjected all 62,753 cells to brain cell-type identification (fig. S6). Here, we adopted a hierarchical clustering strategy (*20, 31*): level-1 clustering classified cells into neuronal cells and glial cells (fig. S7, A and B); level-2 clustering then identified excitatory neurons, inhibitory neurons, astrocytes, oligodendrocytes, oligodendrocyte precursor cells, microglia, and vascular cells (fig. S7, A, C and D); level-3 clustering further identified 57 distinct subtypes (Fig. 3D and fig. S8 to S10). Based on the cell typing results, we generated a spatial cell map from the imaged hemibrain region (Fig. 3E). For example, the spatial cell mapping shows that the cortex is enriched with layer-specific excitatory neurons (Layer 2/3: *Cplx2*^+^, Layer 4/5: *Dkk3*^+^, Layer 6: *Nr4a2*^+^, Layer 6: *Pcp4*^+^), corpus callosum with oligodendrocytes (*Mbp*^+^, *Mal*^+^, *Cnp*^+^, *Plp*1^+^), striatum with medium spiny neurons (*Tac1*^*+*^, *Adora2a*^+^, *Ppp1r1b*^+^, *Penk*^+^, *Rasd2*^+^), thalamus with excitatory neurons (*Prkcd*^+^, *Synpo2*^+^), and hypothalamus with peptidergic neurons (*Hap1*^+^, *Tac1*^+^, *Dlk*^+^) (Fig. 3F). This spatial cell-type map is consistent with previous reports (*31*), demonstrating the potential of RIBOmap single-cell translatomics to comprehensively identify diverse brain cell types and regions. In brief, RIBOmap serves as an alternative and potentially more powerful and accurate strategy for generating a spatial tissue atlas.

In the brain tissue, subcellular localized translation in the cell body (soma) and periphery branches (processes) serves as a critical mechanism to assemble and adjust the network of neuronal and glial cells (*37*–*42*) in response to physiological signals during neurodevelopment and memory. However, it has been a big challenge to analyze mRNA translation within the context of brain networks. Therefore, we set out to test whether RIBOmap further enabled us to study localized mRNA translation directly in intact tissue networks by leveraging the subcellular resolution. To dissect the localized translation in the somata and processes of neuronal and glial cells, we divided the RIBOmap reads into somata-reads (*i*.*e*., inside the cell body area identified by ClusterMap (*43*)) and processes-reads (*i*.*e*., the rest of the reads) (Fig. 4, A to C). Next, we defined the top 10% of genes with the highest and lowest processes-to-somata ratios as processes-and somata-enriched-translation genes, respectively (Fig. 4D). Gene Ontology (GO) analysis showed that processes-enriched-translation genes are associated with translation machinery, synapse, and postsynaptic density (Fig. 4E and fig. S11A). In contrast, somata-enriched-translation genes are associated with the plasma membrane, extracellular matrix, and endoplasmic reticulum (Fig. 4F and fig. S11B), which correspond to localized translated at ER in the somata. Notably, we observed an exceptionally high abundance of processes-enriched-translation signals in hippocampal neuropils, such as postsynaptic density proteins (*e*.*g*., *Shank1, Dlg4, Grik5*), translation machinery protein (*e*.*g*., *Eef2, Eef1a, Rpl3, Rps5*), motor proteins (*e*.*g*., *Kif5a, Kif1a*), and calcium sensor and signaling proteins (*e*.*g*., *Calm1, Camk2n1*) (Fig. 4G and fig. S11C). In contrast, the somata-enriched-translation genes showed sparse RIBOmap signals in hippocampal neuropils (Fig. 4H and fig. S11D), such as *App* and *Rtn4*, which may be associated with ER translation. Besides neuronal genes, we also observed the localized translation in glial cells (Fig. 4, I and J and fig. S11, E and F). For oligodendrocytes, the marker genes *Mbp* and *Plekhb1* were identified as processes-enriched-translation genes, in contrast, *Mal* and *Cnp* were somata-enriched-translation genes (Fig. 4, I and J and fig. S11, E and F). Astrocytes have many processes and their marker genes were enriched in the processes-enriched-translation category as expected, like *Gfap, Mt2, Apoe*, and *Clu* (Fig. 4I and fig. S11E). Vascular cells have very few processes and their marker genes are mostly enriched in the somata-enriched-translation gene group, like *Ptgds, Itm2a*, and *Bsg* (fig. S11F). Overall, we demonstrated that RIBOmap is a powerful tool to study subcellular-localized translation in processes of both neuronal and glial cells of the mouse brain tissue.

**Fig. 4.**
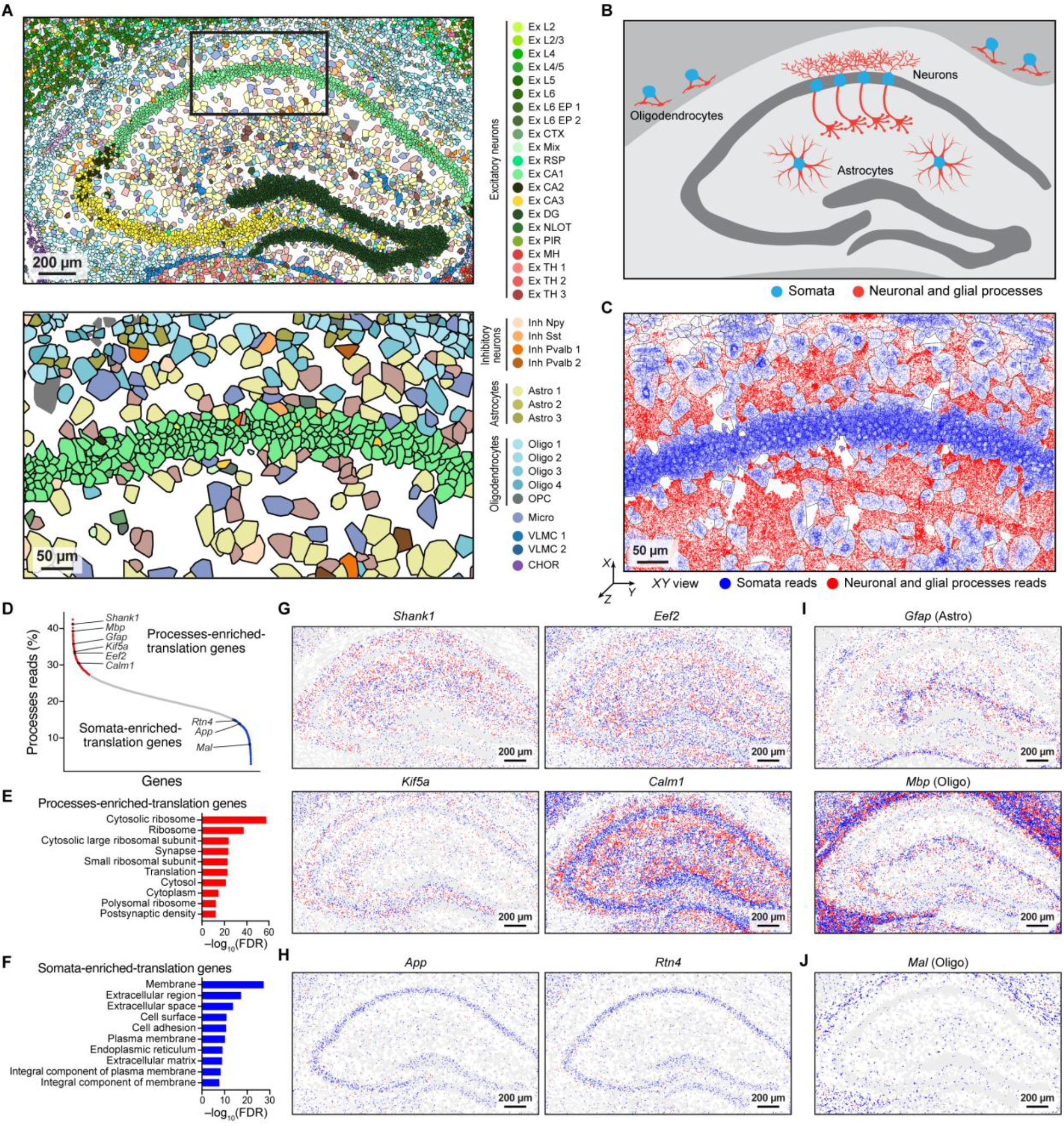
Localized translation in the somata and processes of neuronal and glial cells in the mouse brain. (**A**) Magnified sections in the hippocampus and CA1 region showing the different cell types and the interspace. (**B**) Schematic of a hippocampal slice showing the somata and processes of hippocampal neurons, oligodendrocytes, and astrocytes. (**C**) Section in the CA1 region showing somata reads (blue) and neuronal and glial processes reads (red). (**D**) Processes read percentages of individual genes in the 5,413-gene RIBOmap measurements, with genes rank-ordered based on their processes reads percentage. Nine example genes were labeled inset. (**E)** The top 10 significantly enriched GO terms for processes-enriched-translation genes. (**F**) The top 10 significantly enriched GO terms for somata-enriched-translation genes. GO was analyzed by DAVID. (**G** and **H**) The spatial translation map of representative processes-enriched-translation genes (G) and somata-enriched-translation genes (H) in the hippocampus region, showing somata reads (blue) and processes reads (red). (**I** and **J**) The spatial translation map of glial cell marker gene examples of processes-enriched-translation genes (I) and somata-enriched-translation genes in the hippocampus region, showing somata reads (blue) and processes reads (red).

In summary, RIBOmap is a novel spatial translatomics method with the single-cell and molecular resolutions, opening up a critical step toward a more comprehensive understanding of mRNA regulation and protein synthesis in intact cellular and tissue networks. By examining the mRNA translation during the cell cycle progression, RIBOmap reveals the distinct subcellular distribution patterns of translating RNAs in human cells. Moreover, RIBOmap has achieved the spatial profiling of translatomic states *in situ* in intact tissue samples with spatially resolved and molecular resolution in 3D, generating a translatome-defined spatial cell map of mouse brain tissues. Importantly, in contrast to existing approaches, RIBOmap bypasses complicated polysome isolation steps and genetic manipulation, thus holding great promise for spatial and single-cell resolved studies in *post hoc* human tissue and disease samples. Although RIBOmap in this manuscript focuses on mapping RNA translation, such proximity-based tri-probe design can also be readily adapted to study RNA-RNA interactions, RNA-protein interactions, and RNA modifications. In the future, we envision that RIBOmap can be combined with other imaging-based measurements to enable spatial multiomics mapping of epigenome, transcriptome, and translatome in the same samples for an integrative understanding of biological systems.

## Acknowledgments

We thank Shiyou Zhu (Broad Institute), Ming Pan (Broad Institute) for technical assistance, Haowen Zhou (Broad Institute), Jiakun Tian (Broad Institute), Han Xu (Peking University), Bo He (Peking University), Yafei Yin (Zhejiang University), Jia Liu (Harvard University), Jennifer Lo (Broad Institute) and Leslie Gaffney (Broad Institute) for helpful discussions and thoughtful comments on the manuscript. We thank Feng Zhang (Broad Institute and MIT) for kindly providing the psPAX2 and VSVG plasmids. X.W. acknowledges the support from the Searle Scholars Program, Thomas D. and Virginia W. Cabot Professorship, Edward Scolnick Professorship, Ono Pharma Breakthrough Science Initiative Award, Merkin Institute Fellowship, and NIH DP2 New Innovator Award. H.S. is supported by the Helen Hay Whitney Foundation Postdoctoral Fellowship.

## Funding

The Searle Scholars Program (XW)

Thomas D. and Virginia W. Cabot Professorship (XW) Edward Scolnick Professorship (XW)

Ono Pharma Breakthrough Science Initiative Award (XW) Merkin Institute Fellowship (XW)

National Institutes of Health DP2 New Innovator Award 1DP2GM146245-01 (XW)

Helen Hay Whitney Foundation Postdoctoral Fellowship (HS)

## Competing interests

X.W., H.Z., and J.R. are inventors on pending patent applications related to RIBOmap. All methods, protocols, and sequences are freely available to nonprofit institutions and investigators.

**Fig. S1.**
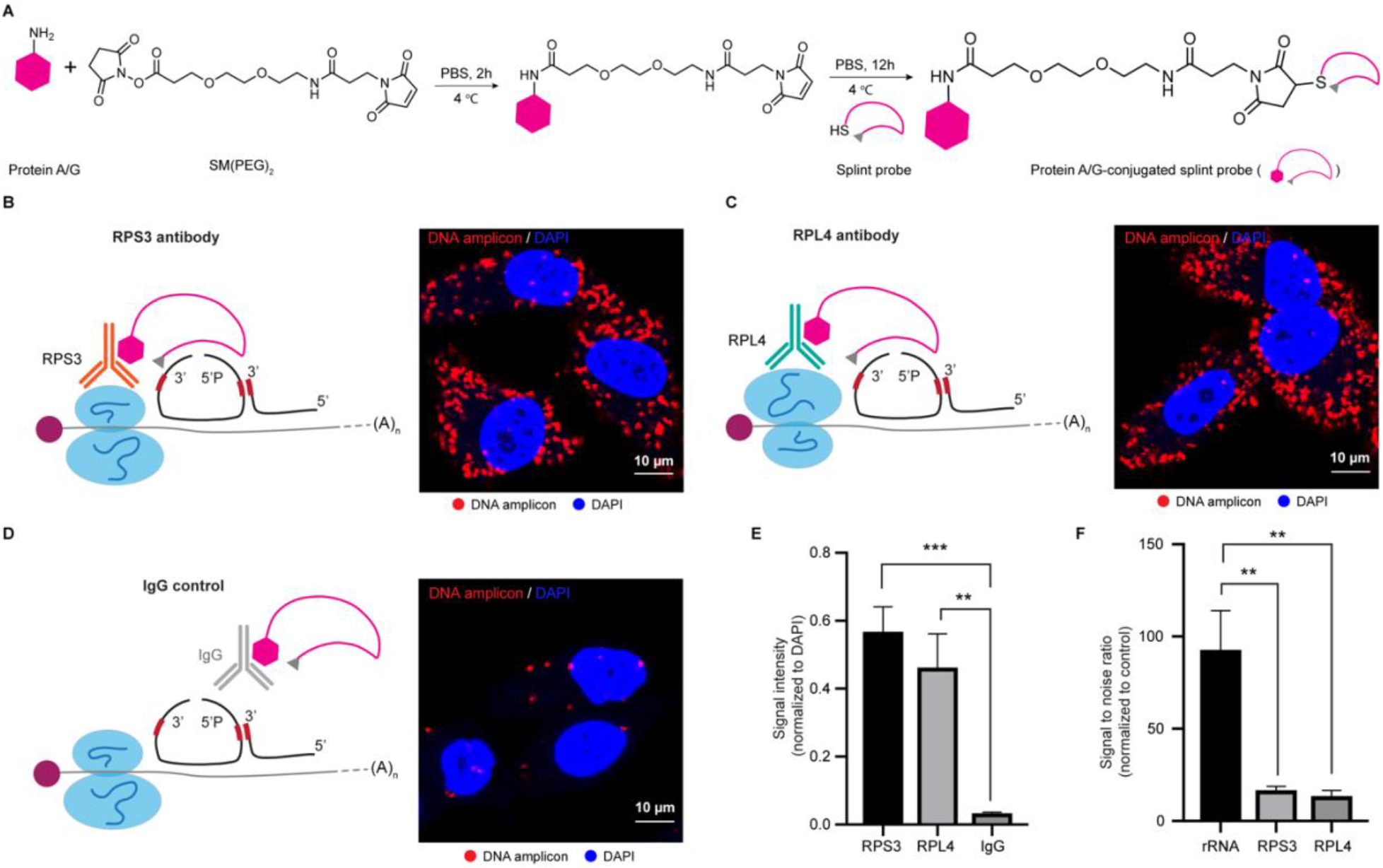
Antibody-based RIBOmap strategy. (**A**) Synthesis scheme for protein A/G-conjugated splint probe preparation. (**B**) Anti-ribosomal protein S3 (RPS3) antibody-assisted RIBOmap signal of *ACTB* mRNA in HeLa cells. RPS3 antibody binds to RPS3 protein in the small unit of the ribosome and brings in protein A/G-conjugated splint probe to the translating mRNA. The padlock probe targeted to the translating mRNA can then be ligated and amplified to generate amine-modified DNA amplicons *in situ*. (**C**) Anti-ribosomal protein L4 (RPL4) antibody-assisted RIBOmap signal of *ACTB* mRNA in HeLa cells. (**D**) The IgG control of the antibody-assisted RIBOmap signal of *ACTB* mRNA in HeLa cells. (**E**) Quantification of the antibody-assisted RIBOmap signal intensity that normalized to DAPI in B-D. Error bars, standard deviation. *n* = 3 images per condition. Student’s t-test, \*\**P* < 0.01, \*\*\**P* < 0.001. (**F**) The quantified comparison of the signal-to-noise ratio of rRNA probe-based RIBOmap strategy and antibody-based RIBOmap strategy. Error bars, standard deviation. *n* = 3 images per condition. Student’s t-test, \*\**P* < 0.01.

**Fig. S2.**
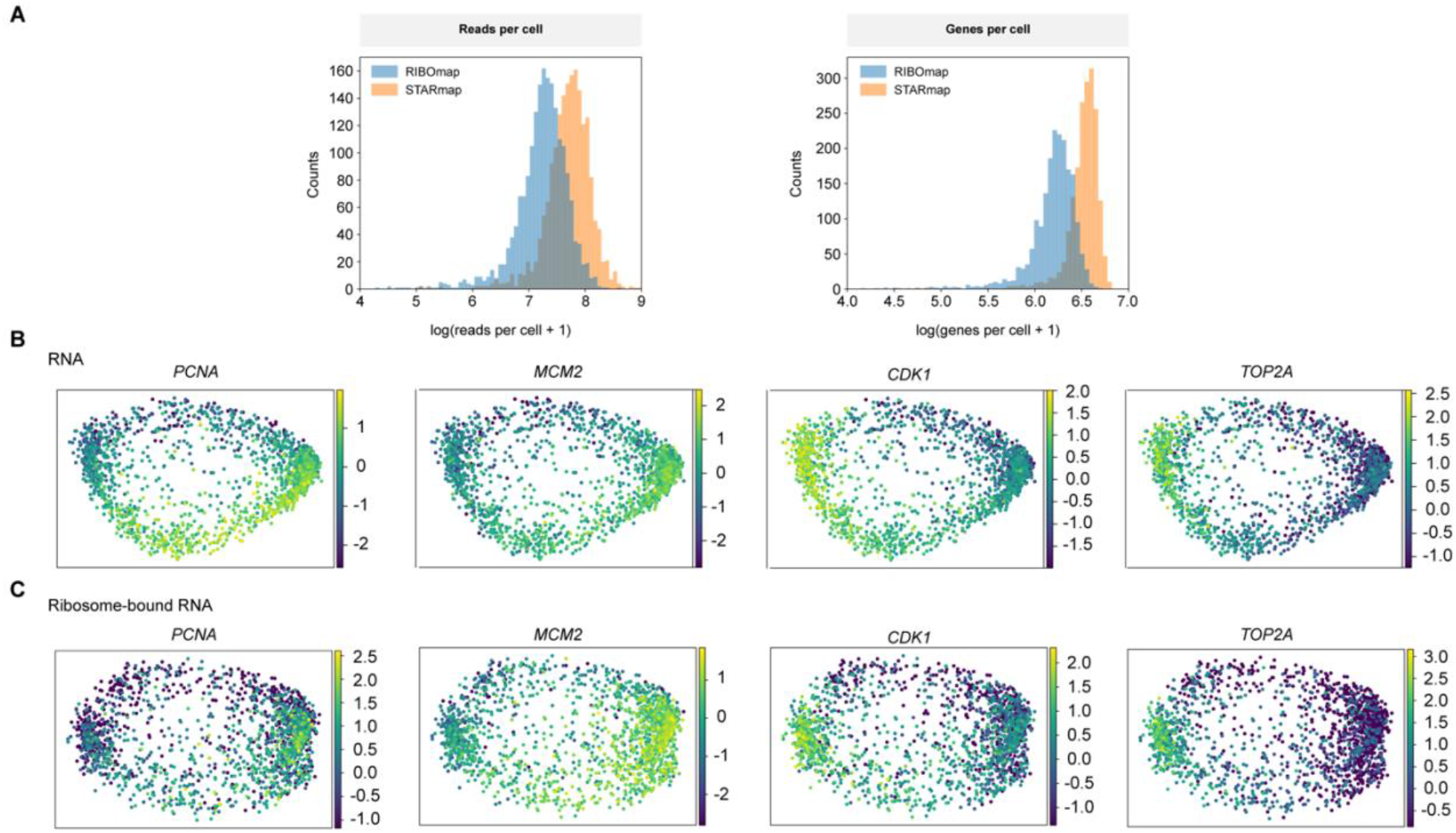
Transcriptomic and translatomic dissection of cell-cycle-related variations. (**A**) Histogram shows the number of transcripts and genes in each cell after logarithmic transformation of the RIBOmap and STARmap HeLa cell dataset. (**B** and **C**) Cell-cycle marker genes expression on diffusion map embedding for cell-cycle stage in STARmap (B) and RIBOmap (C).

**Fig. S3.**
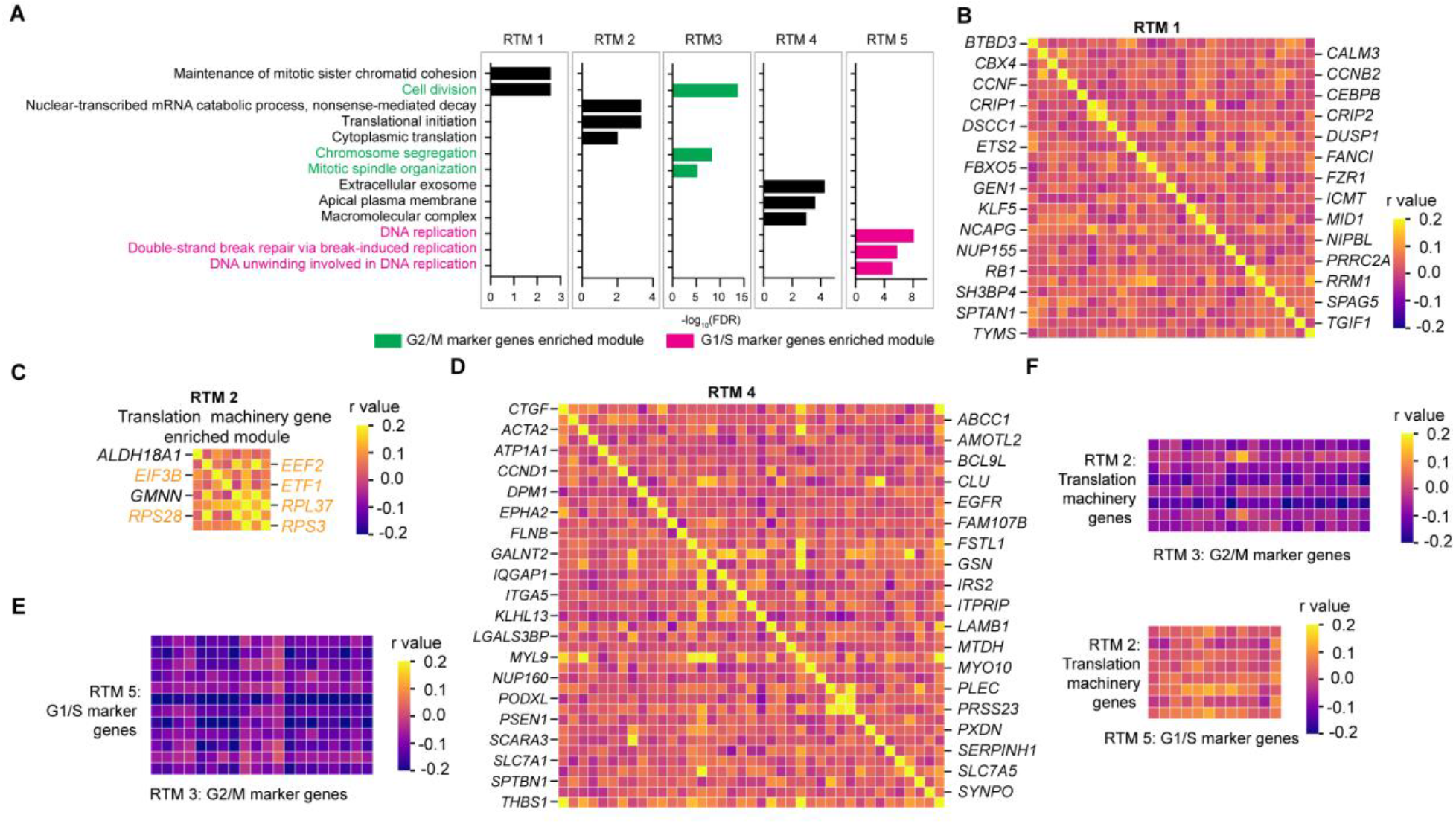
The covariation analysis of single-cell translatome. (**A**) The top three significantly enriched GO terms in each of the five co-regulated translation modules (RTMs) identified in the single-cell translatome covariation matrix. (**B** to **D**) Enlarged RTM 1 (B), RTM 2 (C) and RTM 4 (D) identified in the single-cell translatome covariation matrix. (**E**) Zoom-in view showing the correlation between gene RTM 3 (G2/M marker genes) and gene RTM 5 (G1/S marker genes). (**F**) Zoom-in view showing the correlation between cell-cycle marker gene modules and translation machinery gene modules.

**Fig. S4.**
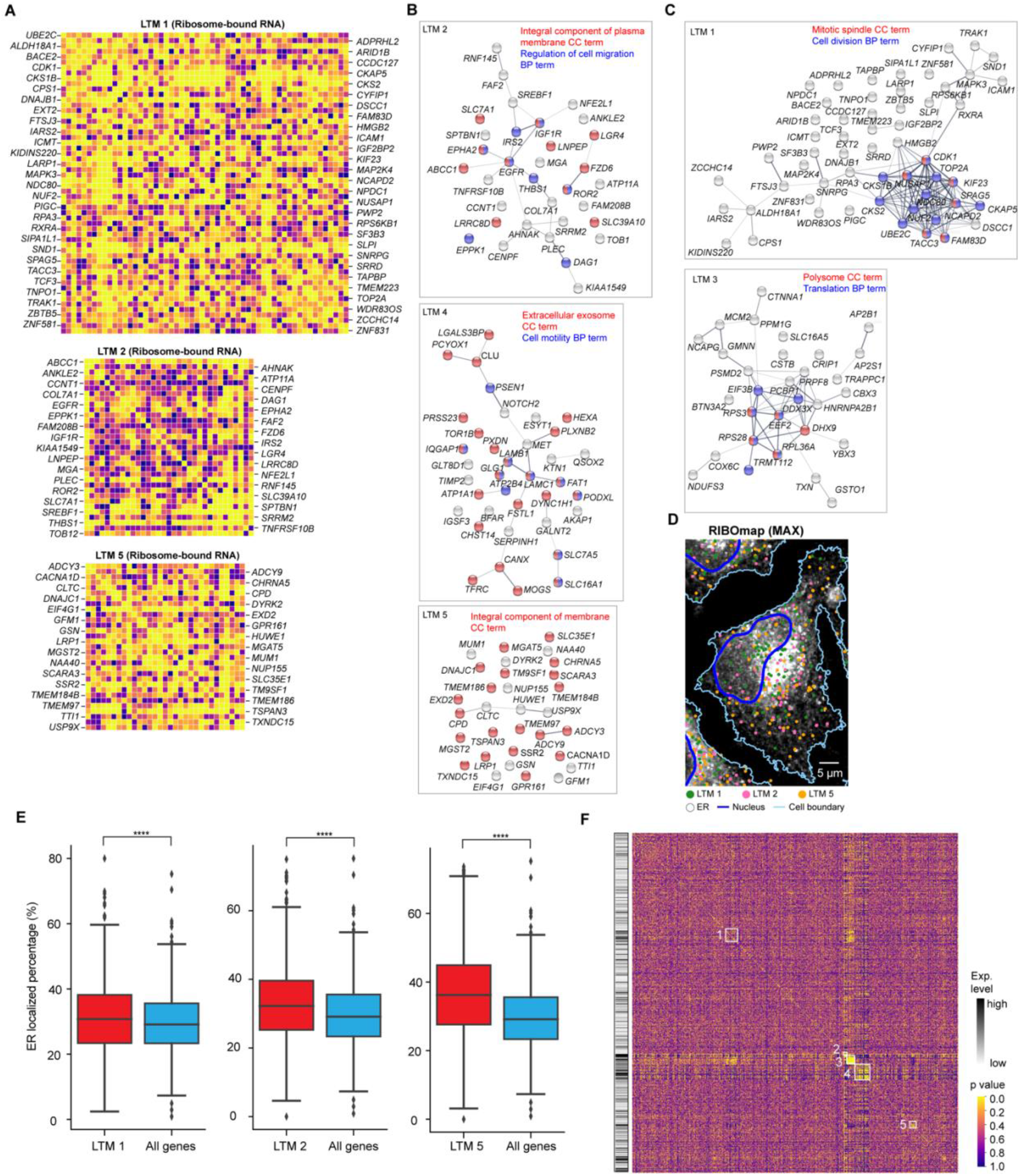
Protein-protein interaction and spatial distribution of co-localized translation modules (LTMs). (**A**) Enlarged LTM 1 (upper), LTM 2 (middle) and LTM 5 (bottom) identified in the ribosome-bound RNA subcellular colocalization matrix. (**B**) The String protein network analysis (*49*) of the genes in LTM 2 (upper), LTM 4 (middle) and LTM 5 (bottom). The enlarged LTM 1 (left), LTM 2 (upright) and LTM 5 (down right). (**C**) The String protein network analysis of the genes in LTM 1 (upper), and LTM 3 (bottom). (**D**) The spatial distribution of RIBOmap signals of the LTM 1, LTM 2 and LTM 5 genes, overlaid on the ER image. (**E**) Quantification of the ER-localized percentage of LTM 1 genes, LTM 2 genes, LTM 5 genes, and all the detected genes. Wilcoxon signed-rank test, \*\*\*\**P* < 0.0001. (**F**) Colocalization pairwise correlation *p*-value on the hierarchical clustering matrix of covariation analysis.

**Fig. S5.**
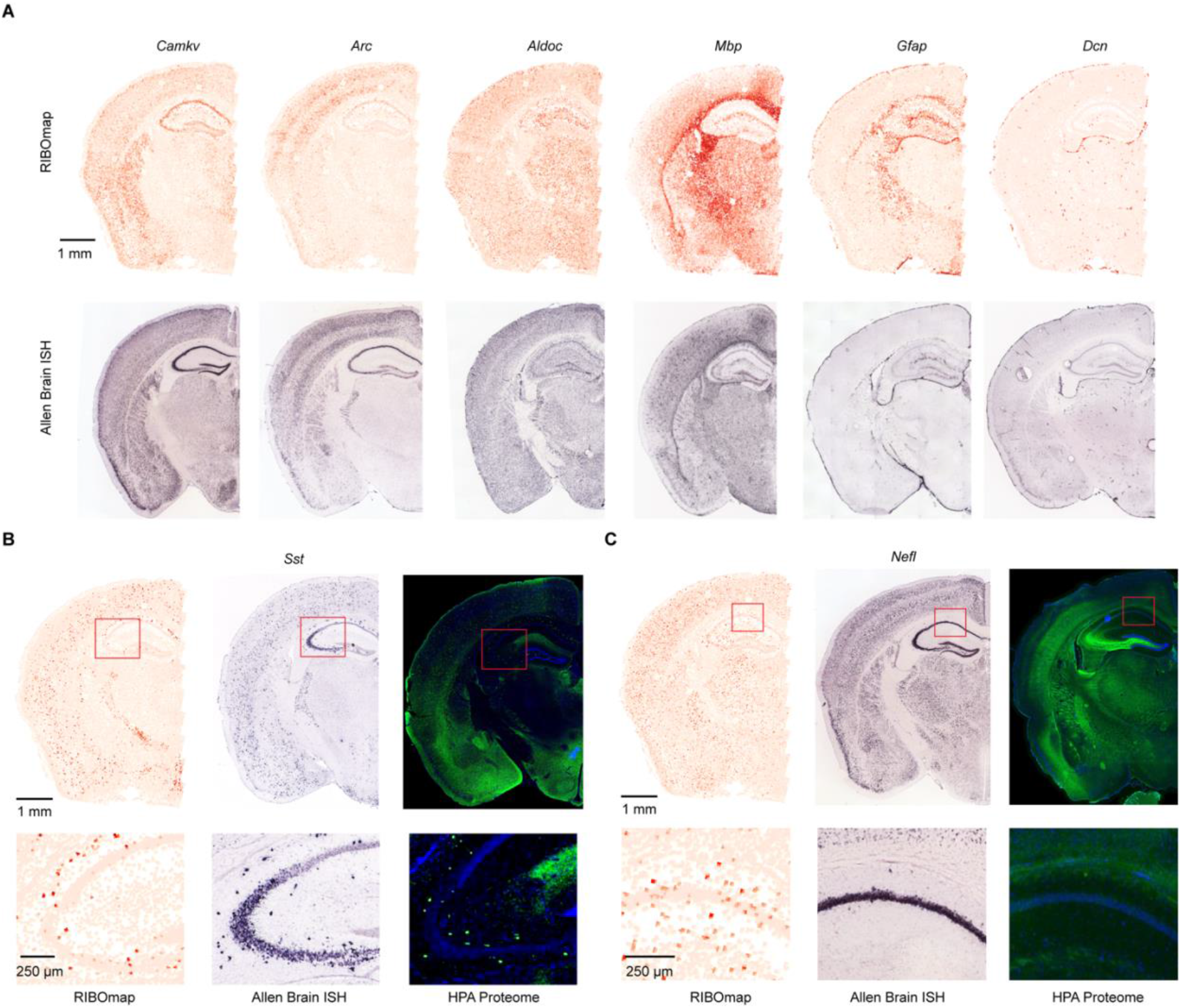
Cross-validation of RIBOmap measurements with in situ hybridization and immunofluorescence. (**A**) RIBOmap images of six example cell marker genes in comparison with the Allen Brain ISH images showing the expression patterns of the corresponding genes. (**B** and **C**) Translation profiles from RIBOmap (this study), *in situ* hybridization (ISH) data from the Allen Brain Atlas and antibody-based immunofluorescence (IF) profiling from the HPA Brain Atlas of *Sst* (B) and *Nefl* (C).

**Fig. S6.**
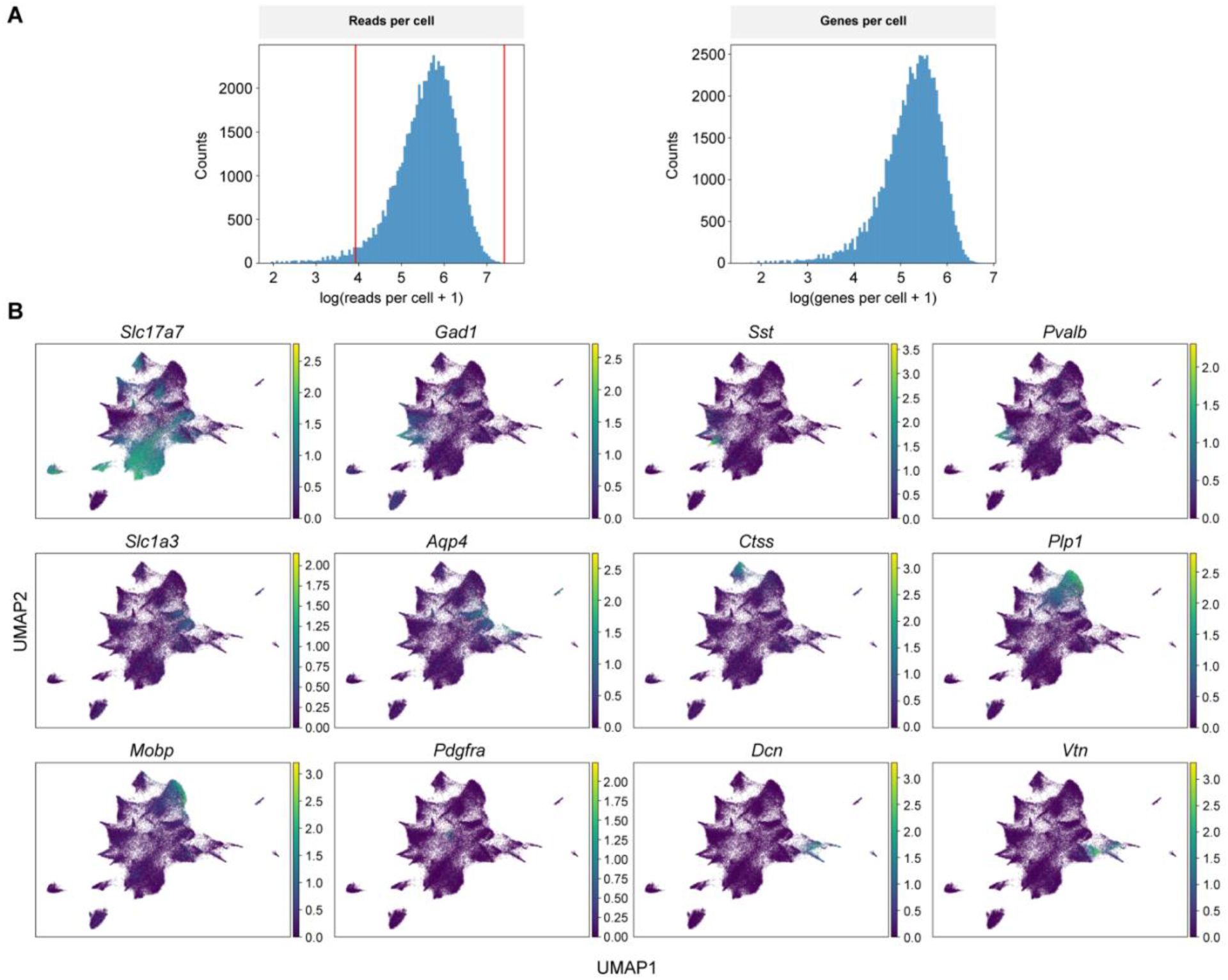
Quality control and cell type marker expression level of RIBOmap mouse brain data. (**A**) Histogram shows the number of transcripts and genes in each cell after logarithmic transformation of the RIBOmap mouse brain dataset. Red vertical lines represent the filtering thresholds estimated by median absolute deviation (MAD). (**B**) UMAPs show expression level of canonical markers of major cell types of mouse brain in RIBOmap dataset.

**Fig. S7.**
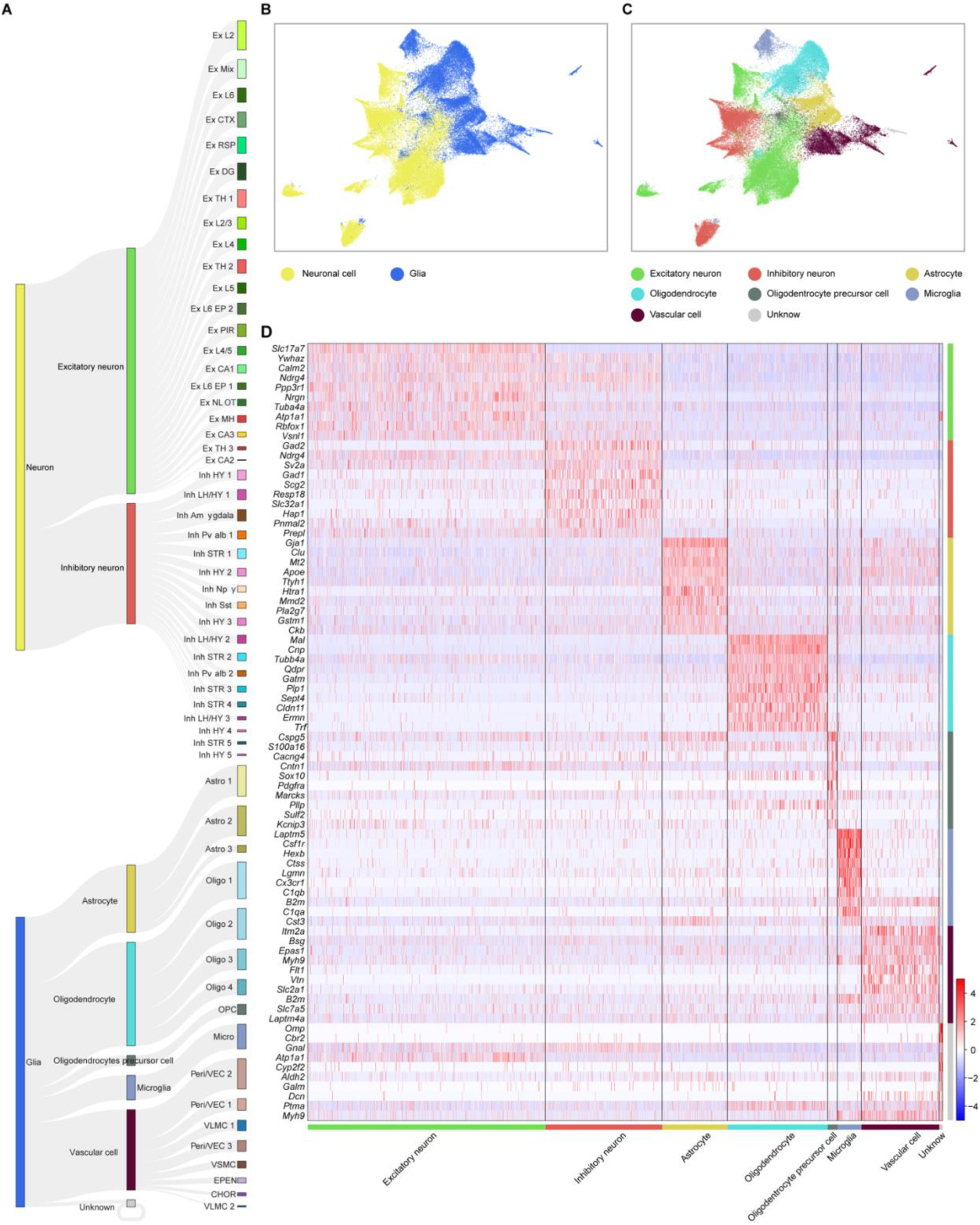
Cell-type clustering based on single-cell translatomics. (**A**) Hierarchical taxonomy of cell types showing the level 1-3 cell-type identification and annotations. (**B**) Uniform Manifold Approximation and Projection (UMAP) plot visualization of the level 1 clustering to classify cells into neuronal cells and glial cells. (**C**) UMAP plot visualization of the level 2 clustering to classify cells into excitatory neurons, inhibitory neurons, astrocytes, oligodendrocytes, oligodendrocyte precursor cells, microglia, and vascular cells. (**D**) Gene expression heatmaps for representative markers aligned with level 2 clustering identified cell types. Expression for each gene is z-scored across all genes in each cell.

**Fig. S8.**
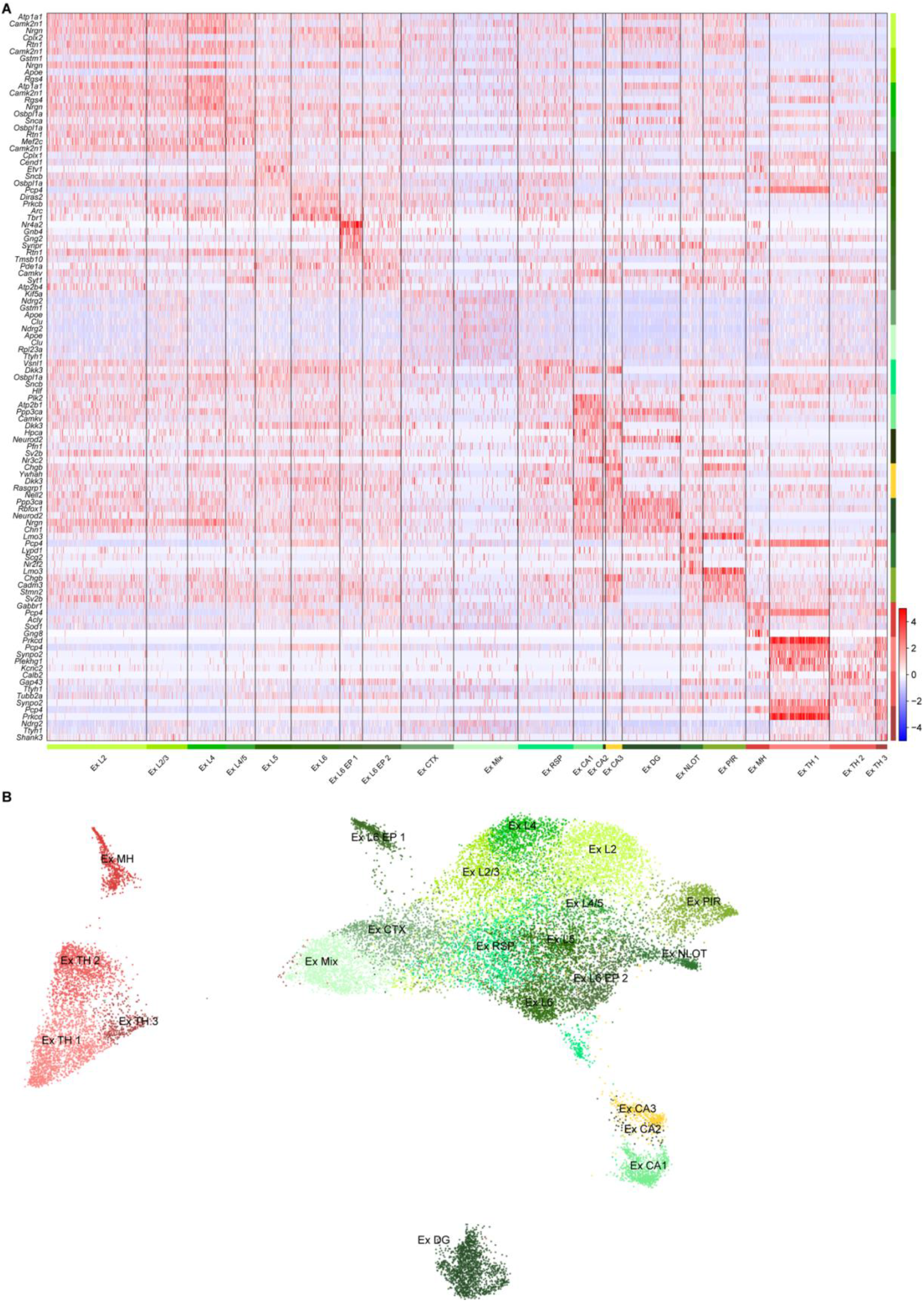
Gene expression and cell clustering of excitatory neurons. (**A**) Gene expression heatmaps for representative markers aligned with excitatory neuron subtypes. Expression for each gene is z-scored across all genes in each cell. (**B**) UMAP plot visualization of the 21 excitatory neuron subtypes.

**Fig. S9.**
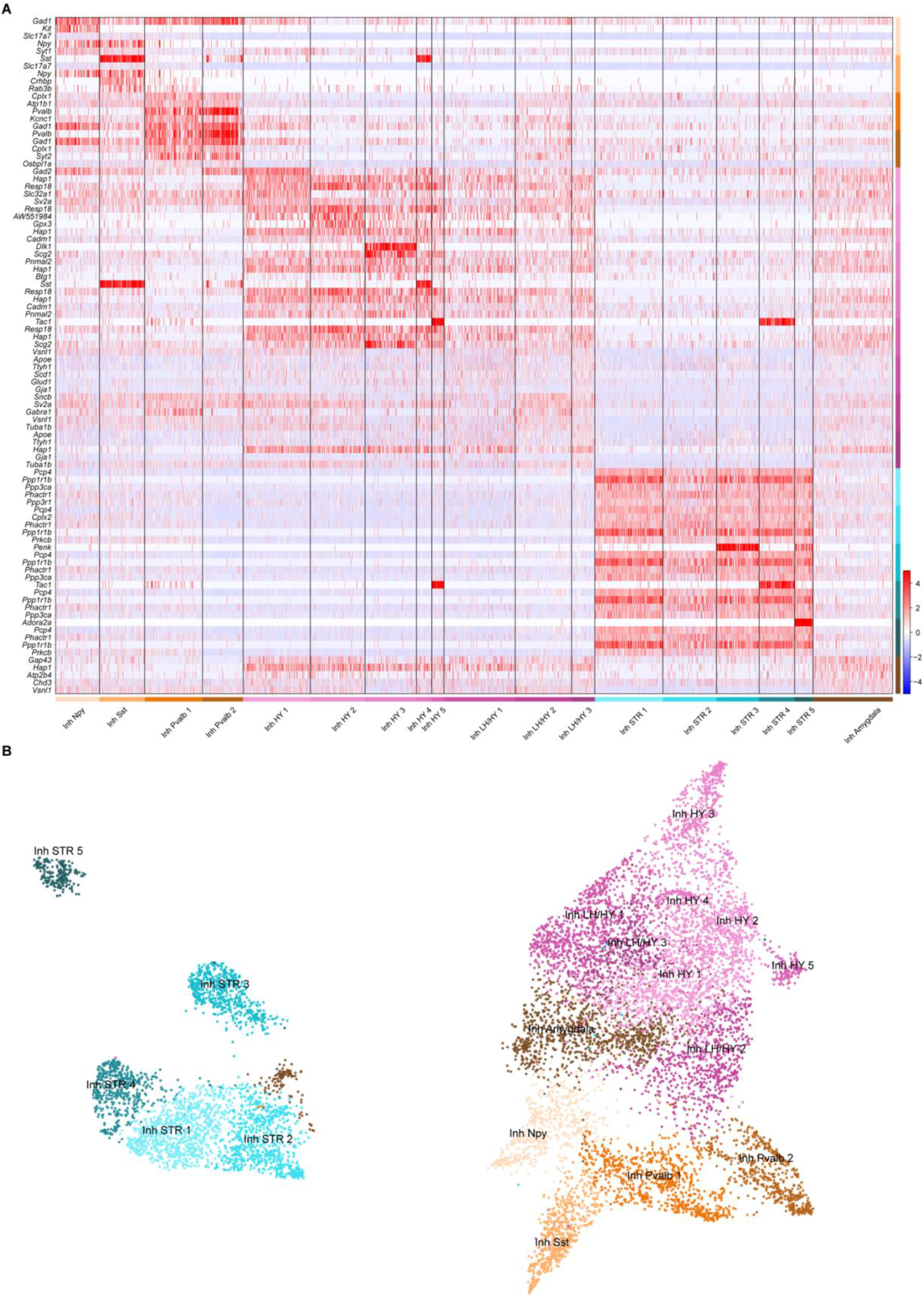
Gene expression and cell clustering of inhibitory neurons. (**A**) Gene expression heatmaps for representative markers aligned with inhibitory neuron subtypes. Expression for each gene is z-scored across all genes in each cell. (**B**) UMAP plot visualization of the 18 inhibitory neuron subtypes.

**Fig. S10.**
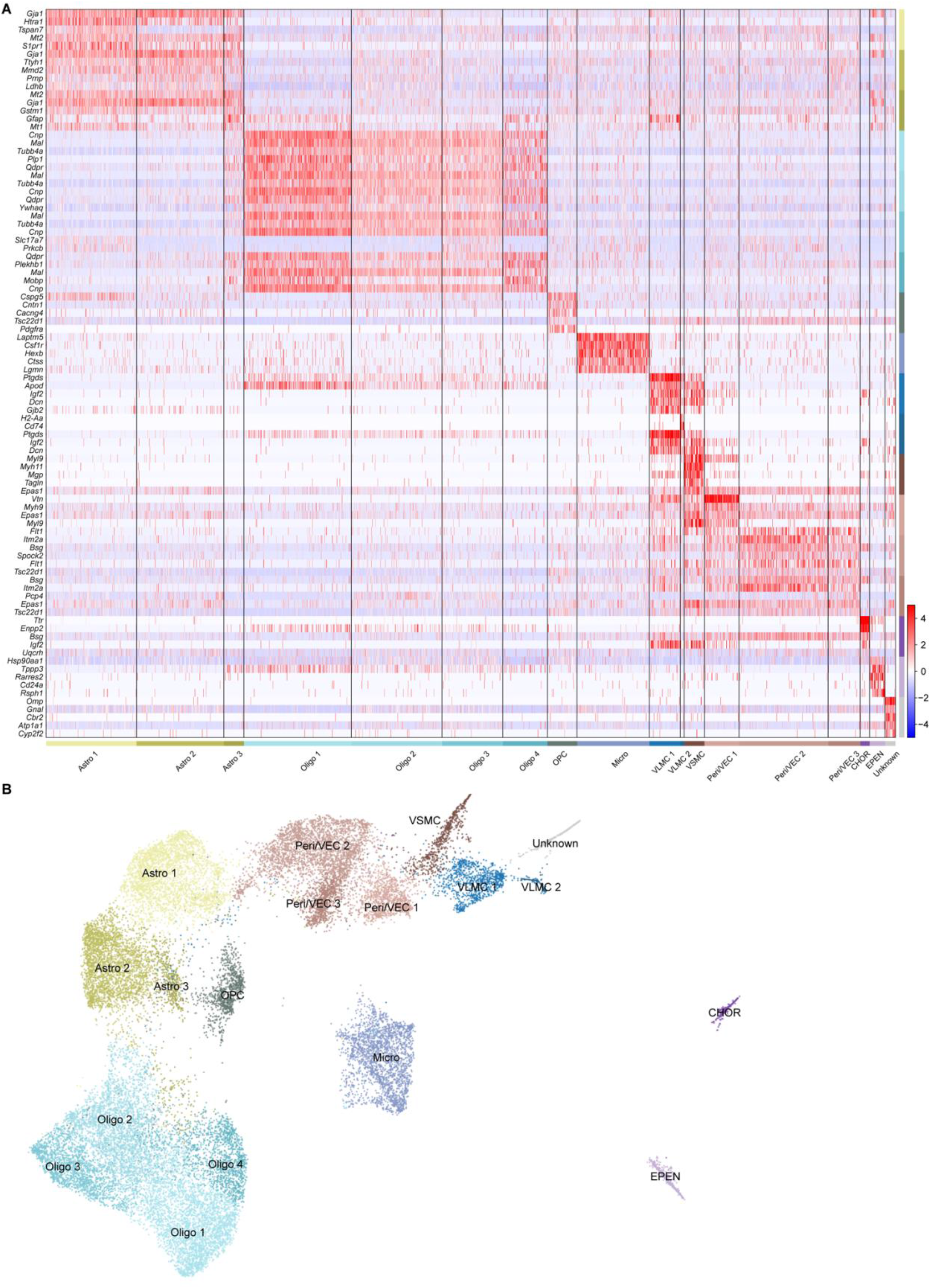
Gene expression and cell clustering of glial cells. (**A**) Gene expression heatmaps for representative markers aligned with glial cell subtypes. Expression for each gene is z-scored across all genes in each cell. (**B**) UMAP plot visualization of the 18 glial cell subtypes.

**Fig. S11.**
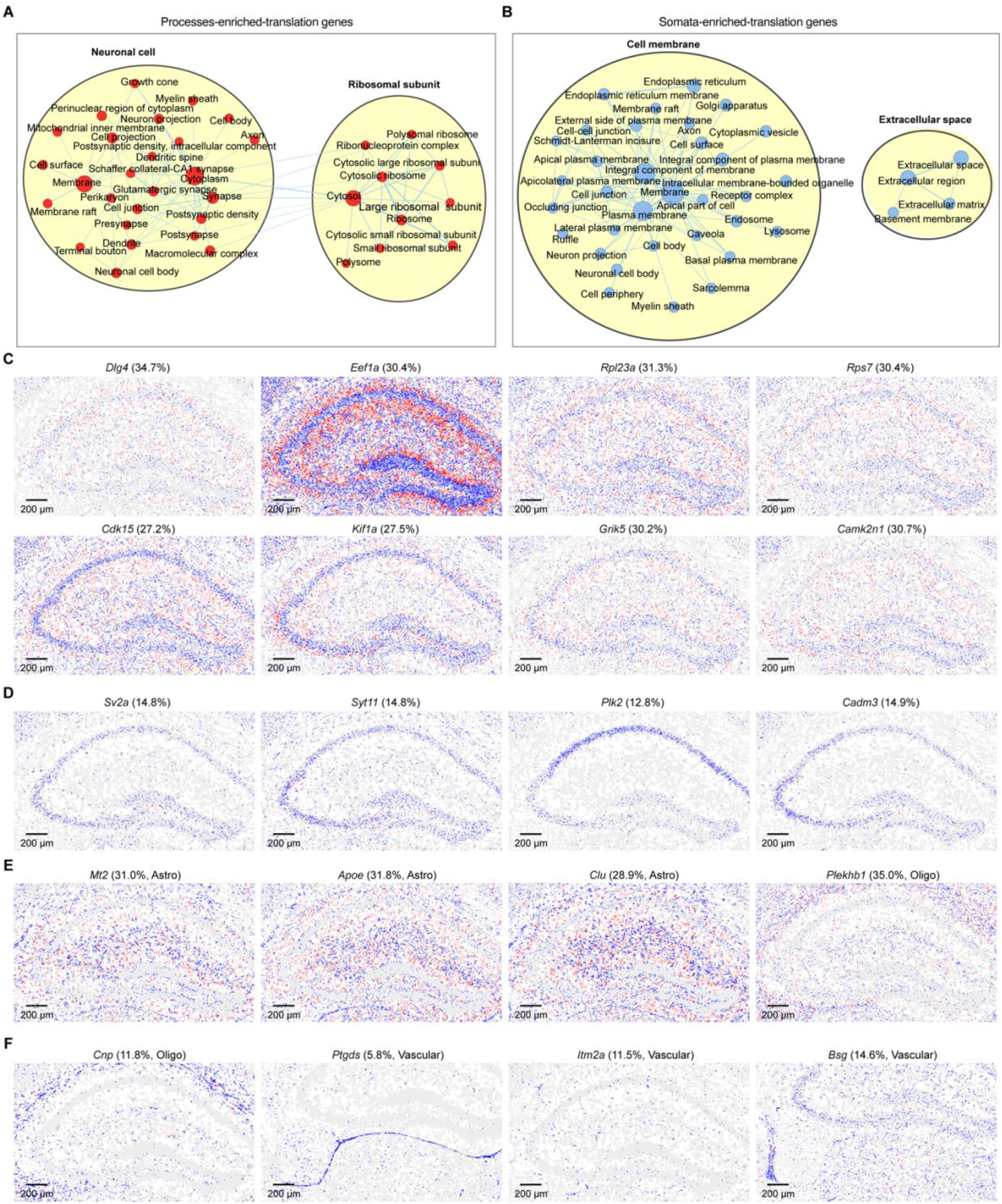
Functional enrichment and examples of processes-enriched-translation genes and processes-enriched-translation genes. (**A** and **B**) A functional enrichment map showing the significant Cellular Component (CC) GO terms of processes-enriched-translation genes (A) and somata-enriched-translation genes (B). (**C** and **D**) The translation spatial map of additional examples of processes-enriched-translation genes (C) and somata-enriched-translation genes (D) in the hippocampus region, shows somata reads (blue) and processes reads (red). (**E** and **F**) The translation spatial map of additional glial cell marker gene examples of processes-enriched-translation genes (F) and somata-enriched-translation genes (F) in the hippocampus region, showing somata reads (blue) and processes reads (red).

